# Dual temporal codes for voice identity in the primate auditory cortex

**DOI:** 10.1101/2025.01.10.632201

**Authors:** Yoan Esposito, Régis Trapeau, Louis-Jean Boë, Luc Renaud, Matthieu Gilson, Thomas Brochier, Margherita Giamundo, Pascal Belin

**Affiliations:** Institut de Neuroscience de la Timone, UMR 7289 Centre National de la Recherche Scientifique & Aix-Marseille Université; GIPSA-Lab CNRS, Université de Grenoble, Grenoble, France; Institute for Language, Communication and the Brain, Aix-Marsielle Université

**Author notes:** co-senior authors (Margherita Giamundo, Pascal Belin).

## Abstract

Recognition of vocal identity is essential for primate social behavior, yet the neural mechanisms supporting this ability remain poorly understood. Cognitive models suggest that the brain may represent identity based on an average voice prototype. Here, we tested this hypothesis by recording spiking activity from single neurons in an fMRI-localized voice-selective region of the anterior temporal lobe in two rhesus macaques. Using synthetic “coo” calls systematically morphed along identity trajectories from “anti-voices” to caricatures, we found that early neuronal responses (∼100 ms post-stimulus) exhibited robust V-shaped tuning to distance from the average voice. This pattern reflects an energy-efficient representational strategy in which neural activity increases with identity distinctiveness, also applied to face stimuli. Unexpectedly, a distinct subpopulation of neurons showed a delayed (∼200 ms) rebound response selectively enhanced for the average voice. These findings reveal two temporally dissociable mechanisms for vocal identity encoding in primate auditory cortex, suggesting that the brain emphasizes both distinctiveness and prototypicality to support efficient and flexible voice recognition.

**Significance statement:** Recognizing who is speaking is fundamental to primate social life, yet how the brain encodes vocal identity remains poorly understood. Cognitive theories suggest that the brain may represent identity relative to an internal “average” voice, but direct neural evidence has been lacking. Here, we recorded single-neuron activity in a voice-selective brain region of rhesus macaques and found two distinct coding strategies: one that increases neural activity with identity distinctiveness, and another that selectively enhances the average voice itself. These results reveal dual mechanisms for vocal identity encoding in the primate brain, highlighting how the brain balances flexibility and efficiency in social communication.

## Introduction

Faces and voices of conspecific individuals are crucial signals in primate social interactions. Over evolutionary time, specialized neural systems have developed to extract and process the rich information these signals convey ^1-3^. Electrophysiological studies in non-human primates, particularly those guided by functional MRI (fMRI), have identified discrete regions in the temporal lobe containing neurons selectively responsive to faces or voices: so-called ‘face cells’ ^4-7^ and ‘voice cells’ ^8,9^. Despite these insights, the coding principles underlying how faces and voices are represented at the neural level remain poorly understood.

A prominent framework in cognitive neuroscience proposes that faces and voices are encoded as locations within a multidimensional feature space, where the center of the space—the “norm”—corresponds to the average face or voice, serving as an identity-neutral prototype ^10-12^. In support of this norm-based coding model, electrophysiological recordings in macaque monkeys have shown that face-selective neurons modulate their firing rates as a function of a stimulus’s distance from the average face ^13-16^. Whether such norm-based principles extend to voice identity coding remains an open question, as direct electrophysiological evidence is currently lacking. However, in support of this notion, fMRI studies in humans have shown that activity in temporal voice areas (TVAs) scales with a voice’s distance to the average: both natural voice variability and morph-based manipulations of identity modulated TVA responses in a distance-dependent fashion ^18^.

In the present study, we addressed this question by recording spiking activity from single neurons in the anterior temporal voice area (aTVA)—an fMRI-defined voice-selective region—in two rhesus macaques. We presented synthetic “coo” calls that were morphed to parametrically vary in acoustic distance from an empirically defined average voice prototype in a multidimensional voice space. Based on the norm-based framework, we hypothesized that neuronal responses would show V-shaped tuning: minimal activity for the average voice, with increasing responses for more distinct identities. Our results confirmed this prediction. Early responses (∼100 ms) exhibited robust V-shaped tuning to distance-from-mean, consistent with norm-based encoding. Unexpectedly, we also observed a delayed (∼200 ms) rebound response in a subset of neurons that selectively enhanced responses to the average voice, revealing a second, prototype-specific coding strategy. These findings suggest that vocal identity is encoded by temporally distinct neural mechanisms—supporting both identity discrimination and prototype representation within primate auditory cortex.

## Results

We analyzed spiking activity from single neurons of two female rhesus monkeys, M1 and M2, implanted with chronic multi-electrode Utah arrays in the fMRI-localized aTVA (Fig. 1A; ^9,19^). The electrophysiological recordings took place while the monkeys performed active detection of a pure tone interspersed amongst a set of synthetic ‘coo’ calls (Fig. 1B). Coo calls are common affiliative vocalizations that allow macaques to recognize the identity of their conspecifics ^20,21^.

**Figure 1.**
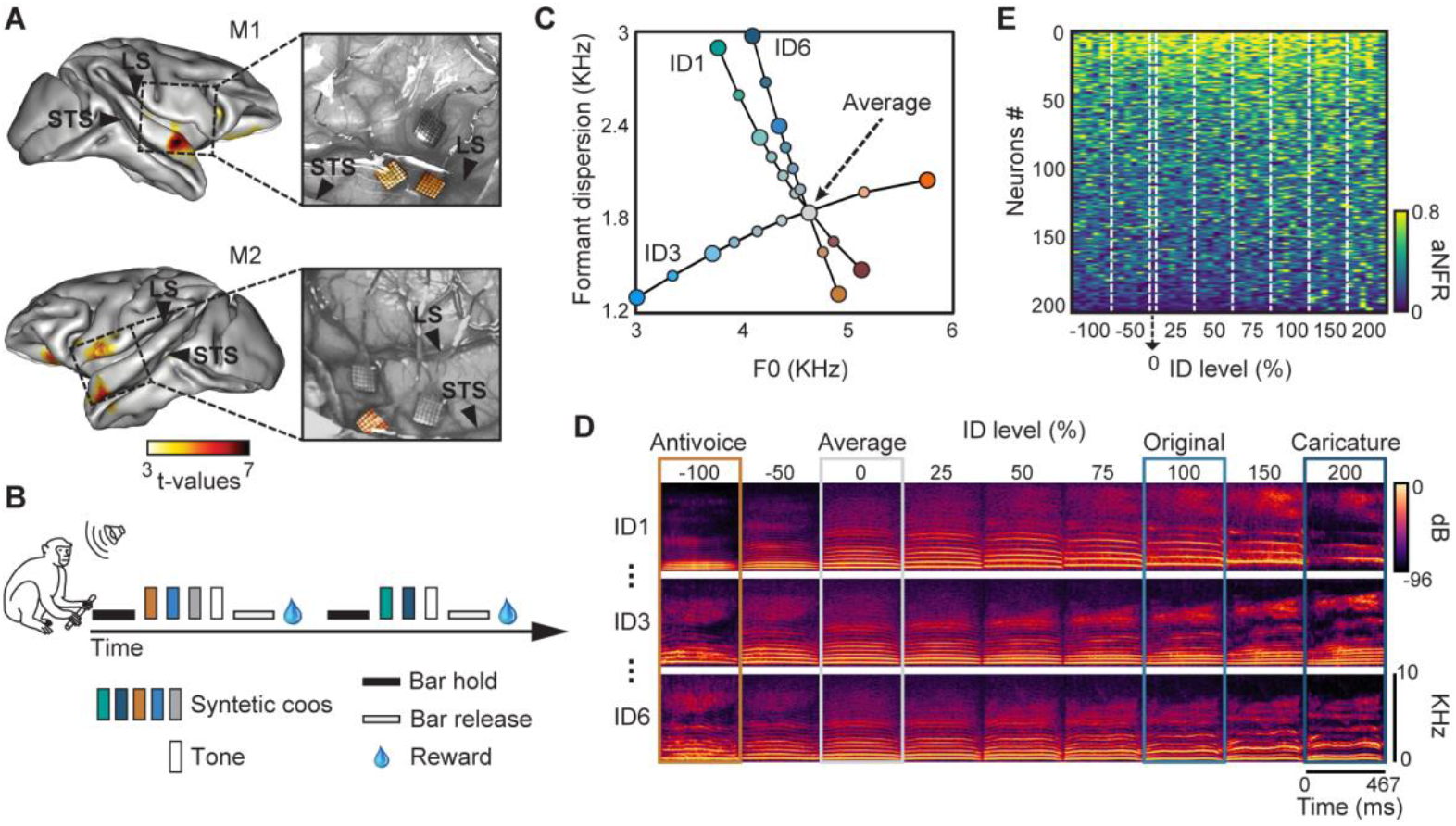
Experimental Design. (**A**) Implantation sites of Utah Arrays, in fMRI defined anterior Temporal Voices Area (aTVA). STS: Superior Temporal Sulcus; LS: Lateral Sulcus. (**B**) The set of 49 vocal stimuli (duration: 467 ms) was presented during a pure tone detection task. In this task, monkeys were required to hold a bar to initiate auditory stimulation and maintain their hold throughout the presentation of vocal stimuli. Only upon hearing a pure tone were they instructed to release the bar in order to receive a juice reward. (**C**) Schematic representation of three example voice trajectories (ID1, ID3 and ID6) in the acoustical voice space. F0: fundamental frequency. (**D**) Spectrograms of three continua generated by morphing between the average and original voices as shown in (C). Identity (ID) levels parametrically represent distance-to-mean, with 0% corresponding to the average coo, 100% the original coo, -100% the anti-voice (opposite acoustical features to those of the original coo relative to the average) and 200% the caricatured coo (exaggerated acoustical differences compared to the average). (**E**) Neurons responsiveness to the 49 stimuli. Each row represents the absolute normalized firing rate (aNFR) of a single neuron, averaged over a 50-150 ms window post-stimulus onset. Neurons are sorted based on their overall aNFR across all stimuli.

Auditory stimuli were generated in four steps (Fig. S1): (1) Recordings of coo calls by 16 different monkeys were first analyzed/resynthesized using STRAIGHTMORPH ^22^. Resynthesized versions of the original recordings were virtually indistinguishable from the originals (at least to a human ear; Supplementary Audio; Fig. S1A). (2) Frequencies of the formants (resonances in the vocal tract) were identified using Linear Predictive Coding, as for human speech (Fig. S1B). (3) Stimuli were morphed together using STRAIGHTMORPH, keeping the first three formants in correspondence across stimuli. Morphing the 16 coos with equal 1:16 weights resulted in a realistic ‘average coo’ with clearly marked first three formants (Fig. S1C). (4) Experimental stimuli were generated with weights corresponding to 6 different ‘identity trajectories’ between 6 of the 16 coos and the average coo (Fig. 1C, D and Fig. S1D; see Methods). Each trajectory spanned morphing steps (‘ID level’) ranging from caricatures (ID level 200% and 150%) to anti-voices (-100%), including the average (0%) and the original stimuli (100%) (Fig. 1C, D). The total stimulus set comprised 49 sounds: 6 identity trajectories × 8 identity levels, plus the average coo (0%) common to all trajectories (see Methods).

We focused our analyses on 205 well-isolated neurons (152 neurons from M1 and 53 neurons from M2) that were significantly modulated by the presentation of a coo call (see Methods; Fig. 1E): 116 neurons (57%) were excited (i.e. with higher spiking rate during stimulus presentation compared to baseline) and 89 (43%) were inhibited.

### aTVA neurons encode distance-to-mean at early latencies

To assess how aTVA neuronal activity was modulated by distance-to-mean, we computed, for each neuron, the average firing rate within a 50–150 ms post-stimulus window, subtracted the baseline activity, and then normalized the resulting values relative to each neuron’s maximal response across the full stimulus set (49 stimuli), yielding a normalized firing rate (NFR). For population-level analyses, we used the absolute value of the NFR (aNFR) to avoid cancellation between excitatory and inhibitory responses.

Figure 2A shows a representative neuron exhibiting robust modulation by distance-to-mean, with a characteristic V-shaped tuning profile with firing rate dropping sharply near the average coo and increasing progressively as stimuli diverged from the average in either direction. This pattern was also evident at the population level (Fig. 2B), as confirmed by a one-way ANOVA across identity levels (0%, -100%, 100%, and 200%) on aNFR (F(3,612) = 17.85, p = 4.05e-11). Post-hoc comparisons revealed significant difference in population responses between the average and both 100% (t = -4.74, p = 2.41e-05) and 200% (t = -5.22, p = 2.70e-06) identity levels, after Bonferroni correction. This norm-based, V-shaped modulation was observed in both monkeys (Fig. S2A).

**Figure 2.**
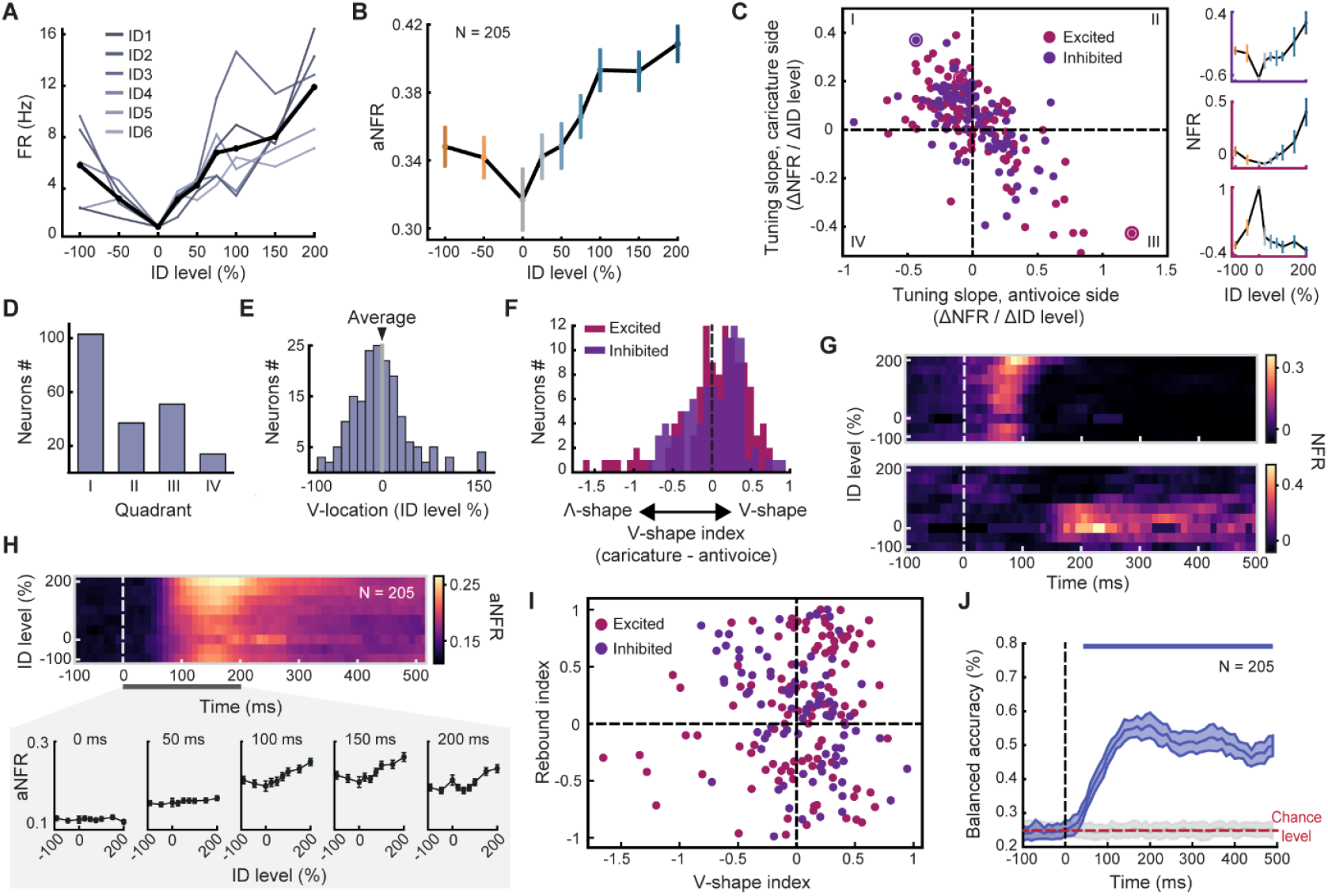
Distance-to-mean differently modulates neural tuning at early and late latencies. (**A**) Example neuron exhibiting a V-shaped tuning. Color lines indicate tuning curves for the different identity trajectories; black line represents tuning curve averaged across identity trajectories. (**B**) Population tuning curve. (**C**) Left panel: scatter plot of anti-voice side slopes vs. caricature side slopes for excited and inhibited neurons. Right panels: Example neurons highlighted in the scatter plot by higher circles. (**D**) Proportion of neurons per quadrants in (C). (**E**) Histogram of identity (ID) levels corresponding to the crossing points of the two regression lines used to estimate the position of each neuron’s response extremum. For illustration purposes, only neurons with crossing points between ID levels -100 and 200% are shown, representing 85% of the total population. (**F**) Distribution of the 205 V-Shape Indices for excited and inhibited neurons. (**G**) Response time courses (heatmaps) of two representative neurons, illustrating distinct temporal dynamics. The top panel shows a neuron exhibiting a typical V-shaped tuning pattern at early latencies, while the bottom panel displays a neuron with a late rebound in activity specifically in response to the average stimulus. (**H**) Top: Population-averaged heatmap showing absolute NFR (aNFR) across identity (ID) levels and time. Bottom: Population tuning curves in 50-ms time bins from 0 to 200 ms post-stimulus onset. (**I**) Scatterplot showing the relationship between the V-Shape Index (capturing early norm-based tuning) and the Rebound Index (capturing late enhancement to the average stimulus) for each neuron. (**J**) Decoding performance (balanced accuracy) of a support vector machine (SVM) classifier trained to discriminate between four identity levels (0, ±50, ±100, and 200%).

To quantify the V-shaped tuning of individual neurons, we performed separate linear regression analyses on NFRs across identity levels ranging from −100 to 0% and from 0 to 200%, following the approach of Koyano et al. (2021). The majority of neurons displayed either V-shaped or inverted V-shaped tuning, reflected by a strong negative correlation between the two slope values (r = -0.67, p = 1.35e-27), confirming sensitivity to distance-to-mean across the population (Fig. 2C, D). To further validate this, we computed the stimulus eliciting the minimal response in each neuron, i.e. the crossing point of the two regressions lines. Consistent with a norm-based coding scheme, 136 out of 205 (66%) neurons had their response minimum within the -50% to +50% identity range, centered around the average stimulus (Fig. 2E).

Finally, we computed a V-Shape Index as the difference between the caricature and anti-voice slopes, where positive values indicate “V” tuning and negative values inverted “V” tuning. A Wilcoxon signed-rank test revealed that the distribution of V-Shape Index values was significantly biased toward positive values (T = 8555, p = 0.019), further supporting the prevalence of norm-based, V-shaped coding in aTVA neurons (Fig. 2F).

### Temporal analysis reveals an activity ‘rebound’ to the average coo at later latencies

To evaluate whether the observed distance-to-mean modulation was stable over time, we performed a temporal analysis of neuronal responses. For each neuron and each identity trajectory, we computed NFR across a sliding temporal window from −100 to 500 ms post-stimulus onset, in 10 ms steps.

Figures 2G and 2H illustrate these temporal dynamics at both the single-neuron (Fig. 2G) and population levels (Fig. 2H). In some neurons, V-shaped distance-to-mean modulation emerged early (around 100 ms; Fig. 2G, top), and was similarly detectable in the overall population around the same latency (Fig. 2H). This early phase reflects a classic norm-based coding profile, consistent with identity tuning relative to the average stimulus.

However, this pattern evolved over time. As shown in Fig. 2G (bottom) and Fig. 2H, around 200 ms post-stimulus onset, we observed a notable rebound in activity that was specific to the average stimulus (identity level: 0%). This late enhancement did not follow the typical V-shape pattern; instead, it reflected a selective increase in response to the average coo, suggesting a distinct, temporally specific process that may represent a different neural mechanism for encoding prototypical identity. The emergence and peak of this rebound (∼200 ms) were consistent across both monkeys (fig. S2B).

To quantify this effect, we developed a “Rebound Index”, measuring the difference in each neuron’s response to the average coo between a late window (150–250 ms) and an early window (50–150 ms). We then compared this Rebound Index to the V-Shape Index, which captures early norm-based tuning. As shown in Fig. 2I, the two indices were not significantly correlated (ρ = –0.002, p = 0.98), suggesting that the late rebound does not emerge from the same neurons involved in early norm-based identity coding. Instead, it appears to be driven by a distinct subpopulation that becomes selectively responsive to the average identity at later latencies.

To further explore the information content of population responses, we conducted a decoding analysis using a linear support vector machine (SVM) classifier. The classifier was trained to discriminate between identity levels 0, ±50, ±100, and 200%, based on population activity. Decoding accuracy rose above chance from 40 ms post-stimulus onset and remained robust throughout the time window (Fig. 2J), confirming that distance-to-mean information is encoded by aTVA neurons in a temporally stable manner.

Lastly, to assess whether distance-to-mean encoding remained stable along time or whether within-session adaptation occurred, we computed the average firing rate of each neuron across identity levels separately for the first and second halves of each recording session. We then generated population tuning curves for both session halves, focusing on two distinct time windows: an early window (50–150 ms post-stimulus), corresponding to the V-shaped tuning, and a later window (150–250 ms), associated with the rebound activity. Our results indicate no evidence of adaptation in either time window for M1, nor in the late window for M2. However, for M2, we observed a small adaptation effect in the early window, specifically for the average stimulus, suggesting a modest within-session modulation of response strength at early latencies (Fig. S3).

### Data-driven decomposition identifies subpopulations with distinct temporal dynamics and distance-to-mean tuning

To ask whether the distinct temporal dynamics we observed for V-Shape and Rebound Indices could be identified in a data-driven manner, we applied Non-Negative Matrix Factorization (NNMF) to the full population activity (Fig. 3A). NNMF is a decomposition method for matrices containing only non-negative values (e.g., spiking rate) widely applied in computational biology ^23,24^. NNMF approximates the non-negative input matrix X as the product of two non-negative matrices: W (weights) x H (factors matrix; Fig. 3A). Using shuffle-split cross-validation, we selected a four-factor solution, as adding more components led to overfitting and reduced generalization performance (Figure S4A). We then used the H matrix to reconstruct the average activity heatmap for each factor. The results revealed that each factor captured distinct temporal and tuning profiles (Figure 3B).

**Figure 3.**
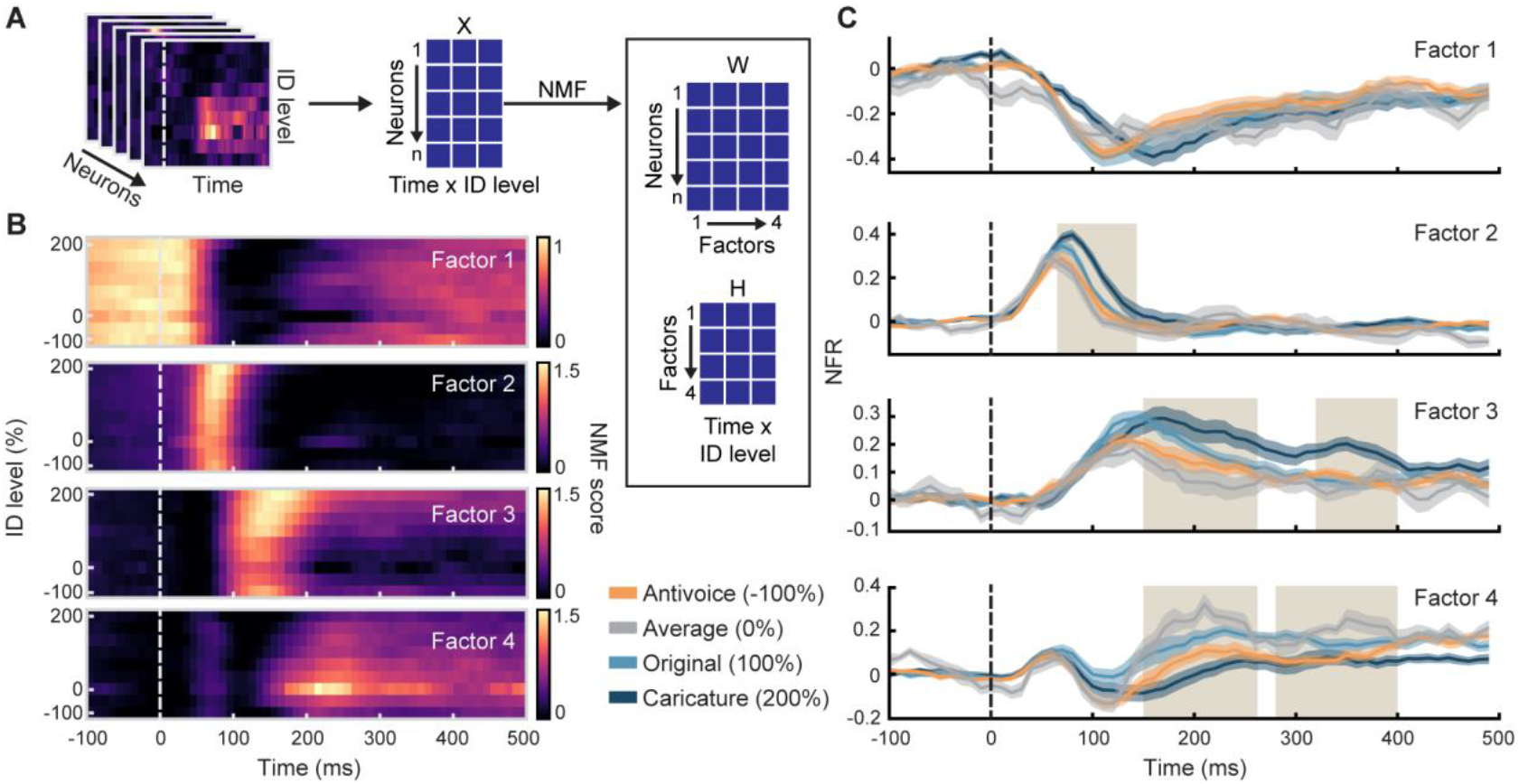
Data-driven Decomposition confirms distinct neuronal subpopulations for Norm-Based Coding and Rebound to the average. (**A**) Non-Negative Matrix Factorization (NNMF) methodology. (**B**) Average heatmap for each of the four factors (top 20% neurons). (**C**) Average population activity of the top 20 neurons of each factor (shaded areas indicate s.e.m.). Vertical shaded areas indicates time bins of significant (p < 0.05, FDR-corrected) distance-to-mean sensitivity.

From the W matrix, we identified the top 20 neurons (top 10%) contributing most strongly to each factor, resulting in distinct neuronal subpopulations. Notably, there was no overlap between these subpopulations — neurons identified as top contributors for one factor did not appear among the top contributors for any other factor (Figure S4B).

To further characterize these subpopulations, we averaged the normalized activity of the neurons within each group, revealing consistent and factor-specific temporal and tuning profiles (Figure 3C). To assess sensitivity to distance-from-mean, we conducted a sliding-window Friedman test across four morph steps (–100, 0, 100, and 200) at each time bin. A subpopulation was considered sensitive to distance-from-mean if the test reached significance (FDR-corrected *p* < 0.05) for at least five consecutive bins.

Factor 1 corresponded to neurons broadly suppressed following stimulus onset, showing no tuning to distance-from-mean. In contrast, the remaining three subpopulations displayed significant distance sensitivity, each with distinct tuning profiles and temporal dynamics. Factor 2 reflected an early V-shaped tuning, with sensitivity emerging rapidly between 70 ms and 140 ms. Factor 3 exhibited a later-onset V tuning, peaking between 150-260 ms and 320-400 ms. Finally, Factor 4 captured a late rebound of activity specific to the average stimulus, with significant sensitivity observed across time windows 150–260 ms and 280–400 ms.

## Discussion

We tested the hypothesis that the spiking activity of voice-patch neurons is modulated by distance-to-mean, following principles similar to those observed in face-patch neurons. To do so, we transposed protocols from the visual domain of face processing ^11,13,16,25,26^ to the auditory modality. Parametric manipulation of distance-to-mean in a set of synthetic coo calls revealed strong modulation of neuronal spiking activity by distance-to-mean, suggesting that the processing of identity is organized very similarly in the face and voice domains.

Particularly, during the early response window (50–150 ms post-stimulus onset), neurons in the aTVA exhibited a V-shaped tuning profile (Fig. 2), consistent with norm-based coding previously described for face patch neurons ^16^. Spiking rates increased with deviation from the average identity, and were further enhanced by caricaturing, suggesting that the neural code in the auditory domain similarly emphasizes identity outliers to facilitate rapid recognition. These findings provide direct evidence that norm-based identity coding extends across sensory modalities, possibly reflecting a general principle of efficient identity encoding in the primate brain.

Despite the overall similarity, three key differences emerged between our results and face patch results:

1. The V-shaped tuning profile was less marked compared to that in face patches. Specifically, whereas in face patch neurons the anti-face elicited as much increase than the original, in our case the increase in the anti-voice side was less marked (Fig. 2). This may be attributed to differences in the stimuli, particularly the limited number of identities (N = 16) used to construct the voice average. However, a possible explanation is also that some voice-patch neurons may encode identity with a linear tuning profile (as suggested by the subset shown in Fig. 2C) paralleling findings from face studies ^15,27^. Expanding the stimulus set in future studies may better reveal whether linear and V-shaped profiles coexist in the auditory identity space.
2. In contrast to the delayed V-shaped response in face patches (emerging at around 200 ms), the auditory V-shaped tuning emerged as early as 100 ms (Fig. 3). In line with this, NNMF identified a component (Factor 4) showing V-shaped structure even before 100 ms (Fig. 4B). This suggests that voice-patch neurons may contribute earlier in the identity-processing hierarchy than face-patch neurons, reflecting modality-specific timing in person recognition circuits.
3. Perhaps the most striking divergence from face-patch responses was a late rebound at ∼200 ms, observed only for the average coo call. Whereas face-patch neurons typically show a suppression to average faces, we observed a rebound enhancement in the voice domain (Fig. 3). This effect was driven by a distinct subpopulation of neurons, as indicated by the lack of correlation between early V-shaped tuning and late rebound indices. Such a temporally specific response may reflect a feedback mechanism enhancing the salience of prototypical voices, potentially for purposes such as template matching or signalling social relevance in macaques. Similar mechanisms have been proposed in visual processing, where average representations serve as stable references for comparison ^13^ although such a ‘rebound’ has not yet been reported in the visual modality. While reminiscent of the ‘normalizing signal’ (delayed suppression in faces) proposed by Koyano et al. (2021), our findings suggest a parallel but distinct auditory mechanism of delayed enhancement.

Overall, the discovery of distinct temporal dynamics and neuronal subpopulations for voice identity encoding underscores the complexity and flexibility of auditory processing in primates, leaving open the possibility that other sound categories could be encoded similarly. By bridging gaps between auditory and visual research, our findings contribute to a broader understanding of sensory encoding and its role in social communication.

## Methods

### Subjects

Data were recorded from two female rhesus monkeys (*Macaca mulatta*, M1 and M2, aged 7 and 8 years, respectively; weight: 5–6 kg). Animal care, housing, and experimental procedures conformed to the National Institutes of Health’s Guide for the Care and Use of Laboratory Animals and were approved by the Ethical Board of Institut de Neurosciences de la Timone (ref: 2016060618508941).

### Alert Monkey fMRI

The monkeys were first scanned for identifying Temporal Voice Areas (TVAs). All the details about the fMRI procedures are reported in Bodin et al. (2021) – in which the two monkeys were called M2 and M3. Here we give only a brief description of these details. Functional scanning was done using an event-related paradigm with clustered-sparse acquisitions on a 3-Tesla MRI scanner (Prisma, Siemens Healthcare), equipped with an 8-channels surface coil (KU, Leuven). Ferrous oxide contrast agent (monocrystalline iron oxide nanoparticle, MION) was used for all the scanning sessions. Monkeys were trained to stay still in the scanner for a fixed period of 8 seconds to receive the reward. To avoid interference between sound stimulation and scanner noise, the scanner stopped acquisitions such that three repetitions of one of 96 stimuli (inter-stimulus interval of 250 ms) were played on a silent background. The 96 stimuli consisted in brief complex sounds from four main categories: human voices, macaque vocalizations, marmoset vocalizations and non-vocal sounds. Then, MION functional volumes were acquired using EPI sequences (multiband acceleration factor: 2, TR = 0.955 s). The analysis included 67 MION runs of M1 and 64 MION runs of M2. Voice selective areas were identified as those regions responding significantly more to conspecific (macaque) vocalizations versus non vocal sounds (Fig. 1A).

### fMRI-Guided Electrophysiology

Monkeys were chronically implanted with high-density microelectrode arrays (CerePort Utah Array, Blackrock Microsystems, Salt Lake City, UT, USA) to record extracellular activity in the fMRI-localized TVAs. Functional maps projected on the individual anatomical surfaces were used to calculate the exact position of the arrays. M1 was implanted with two 32-channels arrays in the anterior TVA (aTVA) of the right rostral Superior Temporal Gyrus (rSTG; parcellation from the D99 macaque brain template; Ts2 from the AC map macaque brain template) and one 32-channels array in the right frontal cortex. M2 was implanted with three 32-channel arrays in the left rSTG (Ts2), of which one in the aTVA. In this study, we analyzed only data collected from the arrays implanted in the aTVA (i.e., two arrays for M1 and one array for M2; Fig. 1A). Electrical signals were amplified and processed using a RZ2 BioAmp Processor (Tucker-Davis Technologies, Alachua, FL, USA) and sampled at 24414 Hz. Raw data collected during recordings were high-pass filtered (300–5000 Hz), and spike sorting was performed offline using the fully automated algorithm MountainSort v5 (Chung et al., 2017) with the SpikeInterface package (Buccino et al., 2020). This algorithm detects, for each channel, individual clusters are then filtered by applying specific thresholds to their quality parameters to identify single neurons (Chung et al., 2017). Clusters with very low firing rates (FRs) were discarded. Finally, the mean waveforms of the remaining clusters were visually inspected to exclude neurons with irregular shapes.

### Auditory Stimuli

Auditory stimuli were generated using STRAIGHTMORPH (Belin & Kawahara, 2024) in Matlab. We started from recordings of isolated coo vocalizations from 16 different macaques kindly provided by Marc D. Hauser and Asif A. Ghazanfar. Author L.J.B performed LPC-based estimation of formant frequencies for each vocalization. First, an average (0% identity) was generated by morphing 16 original “coo” vocalizations together, using the frequencies of the first three formants as spectrotemporal landmarks put in correspondence across stimuli. Then, six coo continua were generated by morphing between the average and each of six different original coos, resulting in 6 identity trajectories. On each trajectory were generated 9 stimuli: the average (0% identity); the original (100% identity); caricatures (150% and 200%) identity; anti-caricatures (25%, 50% and 75% identity); and ‘anti-voices’ (-50% and -100% identity). The total stimulus set comprised 49 sounds: 6 trajectories × 8 morphing steps, plus the identical average coo (0%) common to all trajectories. All stimuli had a duration of 467ms and were mono, 16bits, with a sampling rate of 22050Hz.

### Experimental Design and Task

All recordings were performed in an acoustically insulated room. The monkeys sat in a primate chair with the head fixed by a non-invasive modular restriction mask (MRM) developed in our laboratory. Auditory stimuli were presented through a RZ6 Multi-I/O Processor (Tucker-Davis Technologies, Alachua, FL, USA) and transduced by two 8020 Genelec speakers, which were positioned at ear level 72 cm from the head and 60 degrees to the left and right. Stimuli were delivered at a sound pressure level of approximately 92 dB. Hand detection was achieved using two optical sensors. Monkeys were trained to perform a pure tone detection task (Fig. 1B). This task was introduced to maintain the attention of the monkeys on the auditory stimulation. They were required to hold a bar with both hands for 1500-2000 ms to trigger the presentation of the sounds. In each trial, from three to seven stimuli (inter-stimulus interval: 280-540 ms) were played after which a 500 ms 1000-Hz pure tone was presented. The pure tone instructed the monkeys to release the bar to receive the juice reward (correct trials). If the monkeys released the bar before the pure tone presentation (false alarm trials) or did not release the bar (miss trials; upper reaction time: 250 ms), no reward was given.

### Data analysis

The analyses were performed using Python software.

#### Auditory responsive neurons

To determine if a cell was auditory responsive, a T-test was performed between baseline period (-100 ms to 0 ms from stimulus onset) and stimulus presentation period (0ms to 500ms from stimulus onset). Cells that exhibited significance (p < 0.05) were further analyzed. Positive and negative t-values were used to classify neuronal activity as either excitatory or inhibitory.

#### Normalization

For neuronal activity normalization, we followed the methodology described in Koyano et al. (2021). Specifically, the response to each stimulus was normalized by subtracting the baseline response and dividing it by the absolute value of the maximum difference to the baseline. This procedure yielded a normalized firing rate (NFR) ranging from –1 to 1, where negative values indicate inhibition and positive values indicate excitation.

#### Population tuning curve

To compute the population tuning curve, average firing rate per ID level was computed and normalized for each cell, as described above. We then averaged normalized activity for each ID level across all neurons.

#### Response time course

Neuronal activity was analyzed using 50 ms sliding windows with a step size of 10 ms, spanning the time interval from –100 ms to 500 ms relative to stimulus onset, resulting in 60 time bins. For each bin, we computed the average firing rate across the 49 stimuli, yielding a matrix of shape [s, *t*], where *s* s is the number of stimuli and *t* the number of time bins. We then applied the same normalization procedure described previously: the baseline response was subtracted from each value in the matrix, which was then divided by the absolute difference between the baseline and the matrix’s maximum value. This produced a normalized firing rate matrix with values ranging from –1 to 1, where negative values reflect inhibition and positive values reflect excitation. We finally averaged normalized activity per ID level, leading to a new matrix of shape [*m, t*], where *m* is the number of ID levels and t the number of time bins. Importantly, each 50 ms window was centered, aligning the normalized firing rate to the midpoint of each bin. This approach enabled a fine-grained examination of the temporal dynamics of neuronal responses.

#### V-tuning modelling

To quantify the proportion of neurons exhibiting “V”-shaped tuning, we adapted framework from Koyano et al. (2021). Specifically, we calculated two regression lines to describe firing rates across ID levels on either side of the average (−100% to 0% identity and 0% to 200% identity). These lines allowed us to determine the slope of the firing rate change per 1% change in ID level.

The V-shape Index of a neuron was defined as the difference between the slope of the regression line on the caricature side and the slope of the regression line on the anti-voice side.

#### Rebound Index

The Rebound Index for each neuron was computed as follow:

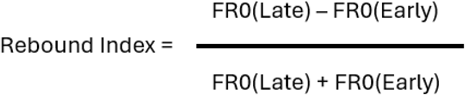

where FR0(Early) is the average firing rate for the average stimulus in the 50-150ms time window from stimulus onset and FR0(Late) is the average firing rate for the average stimulus in the 150-250ms time window from stimulus onset. A Rebound Index of 0.33 indicates that the firing rate of the neuron for the average stimulus has doubled compared to the early window.

#### Decoding Analysis

To investigate the temporal dynamics of neural population coding, we implemented a decoding framework inspired by the Neural Decoding Toolbox (Meyers, 2013). For each decoding run, a pseudo-population response matrix was constructed by sampling trials across recorded neurons. Spike counts were binned in time using a sliding window (e.g., 50 ms bins with a fixed pre-stimulus window), and the response of each neuron in each bin formed the features used for classification. A linear support vector machine (SVM) was trained to predict stimulus ID level (only 0, 50/-50, 100/-100 and 200) based on these binned population vectors. We used a stratified cross-validation procedure (typically 5-fold) to ensure balanced representation of stimulus conditions across training and testing splits. To obtain stable estimates, decoding was repeated multiple times (e.g., 50 runs) with different trials sampled at each iteration. Decoding balanced accuracy was averaged across runs to yield a final performance measure for each time bin. Statistical significance of decoding results was assessed using permutation tests, in which stimulus labels were randomly shuffled (e.g., 100 permutations) to generate a null distribution. Results were considered significantly above chance if they exceeded all values in the null distribution for 5 consecutive time bins (P-value threshold of P = 1/(50 * 60)= 0.0003, corrected for multiple comparisons).

#### Non-Negative Matrix Factorization

We applied Non-Negative Matrix Factorization (NNMF) to explore the decomposition of neuronal population activity. Specifically, we computed the average firing rate of each neuron for each morph step within the time window from –100 ms to 500 ms relative to stimulus onset, resulting in a matrix of shape *m, t+, where *m* is the number of morph steps and *t* the number of time bins. This matrix was then normalized by dividing all values by its maximum, yielding a normalized activity matrix with values ranging from 0 to 1. We performed this normalization independently for each neuron, resulting in a final data structure of shape *n, m, t+, where *n* is the number of neurons, *m* the number of morph steps, and *t* the number of time bins. This matrix was reshaped to [n, m * t] for NNMF computation. To identify the optimal number of factors, we employed a repeated Shuffle Split cross-validation procedure (n = 50 splits, test size = 50%), assessing the stability and generalizability of the NNMF decomposition. For each value of *k* (1 to 10 factors), the NNMF model was trained on the training data and subsequently used to transform and reconstruct the held-out test set and the explained variance between the original matrix and the reconstructed one was then computed. The factorization matrix was finally used to reconstruct the average heatmap for each component.

## Funding

Fondation pour la Recherche Medicale AJE201214 (PB)

Agence Nationale de la Recherche ANR-16-CE37-0011-01 (PRIMAVOICE)

Agence Nationale de la Recherche ANR-16-CONV-0002 (Institute for Language, Communication and the Brain) (MG, PB)

Agence Nationale de la Recherche ANR-11-LABX-0036 (Brain and Language Research Institute) (PB)

Excellence Initiative of Aix-Marseille University (A∗MIDEX) (PB)

European Research Council (ERC) under the European Union’s Horizon 2020 research and innovation program (grant agreement no. 788240) (PB)

## Author contributions

Conceptualization: RT, MG (Margherita Giamundo), PB

Methodology: RT, LJB, LR, TB, MG (Margherita Giamundo), PB

Software: RT

Investigation: MG (Margherita Giamundo)

Resources: PB

Data curation: RT, MG (Margherita Giamundo)

Formal analysis: YE, MG (Matthieu Gilson)

Writing – original draft: YE, MG (Margherita Giamundo), PB

Writing – review & editing: YE, RT, LJB, LR, MG (Matthieu Gilson), TB, MG (Margherita Giamundo), PB

Visualization: MG (Margherita Giamundo)

Supervision: MG (Margherita Giamundo), PB

Project administration: MG (Margherita Giamundo),

PB Funding acquisition: PB

## Data and materials availability

The preprocessed data and relevant metadata in this paper will be available in the Zenodo repository.

## Supplementary Materials

Fig. S1 to S4

Audio S1

## Supplementary Figures

**Fig. S1.**
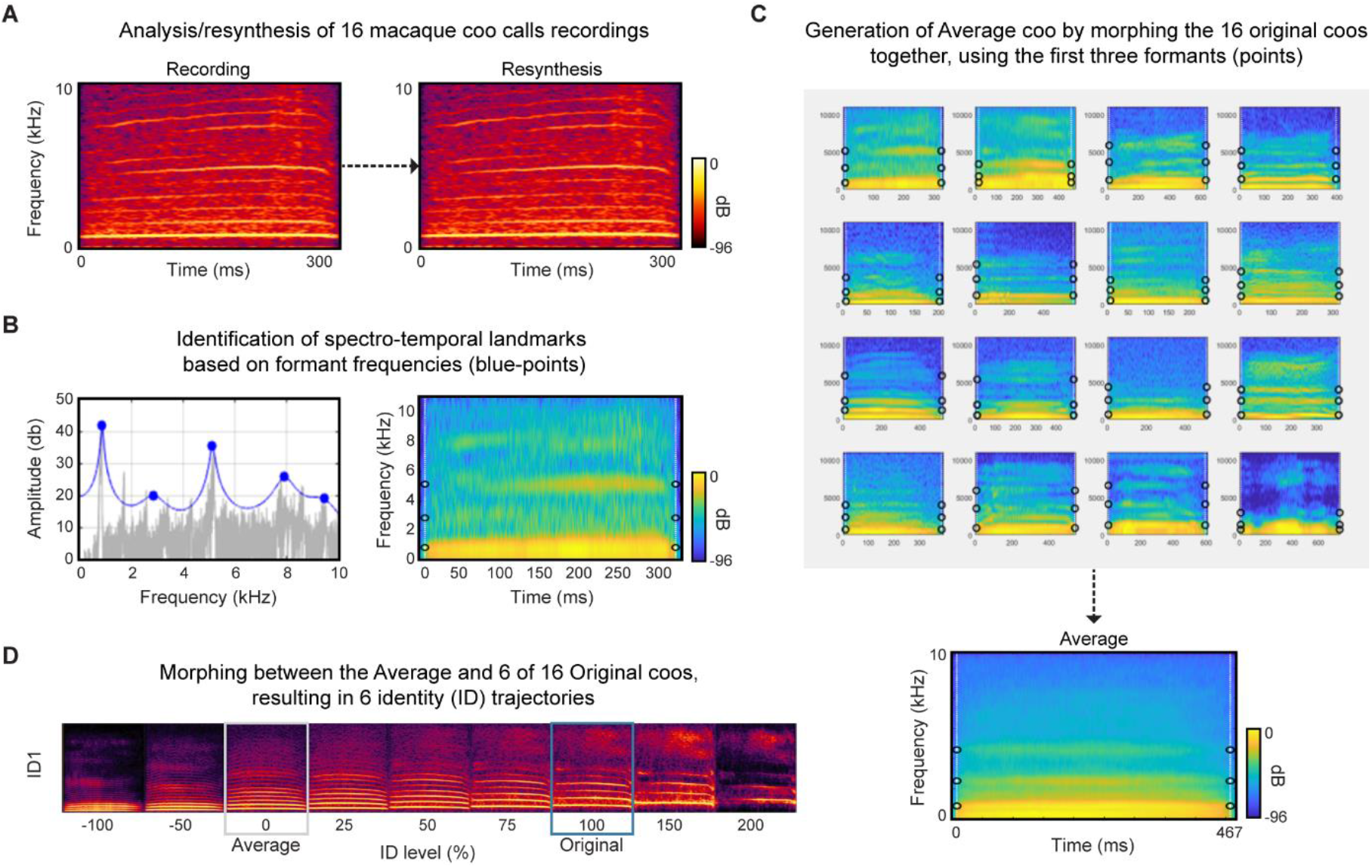
Stimulus generation. **(A)** Recordings of ‘coo’ vocalizations from 16 monkeys were resynthesized using STRAIGHTMORPH ^22^; (B) Formant frequencies were identified using Liner Predictive Coding; (C) The 16 coos were morphed together, using the frequencies of the first 3 formants as spectrotemoral anchor points put in correspondence across stimuli, yielding e.g. to 16-coo average (bottom panel); (D) Distance-to-mean Continua were generated by morphiong between one of 6 Original coos and the average coo, form caricature to ‘anti-voices’. **Cf. Suppl. Audio 1**

**Fig. S2.**
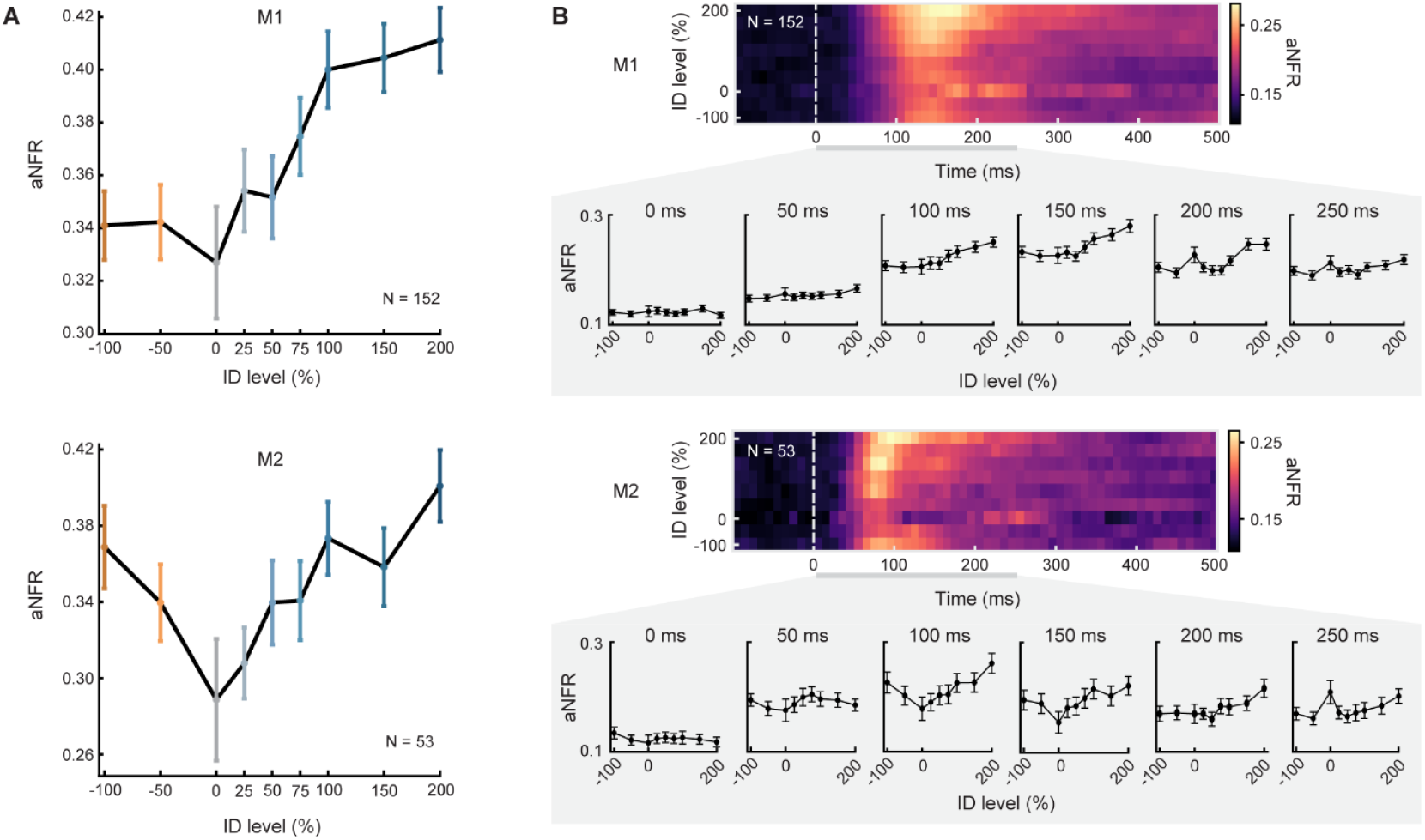
Both monkey exhibit similar tuning and temporal dynamics. (**A**) Tuning curves for each monkey at early latencies (aNFR, 50-150ms from stimulus onset). (**B**) Population heatmaps for each monkey (aNFR).

**Fig. S3.**
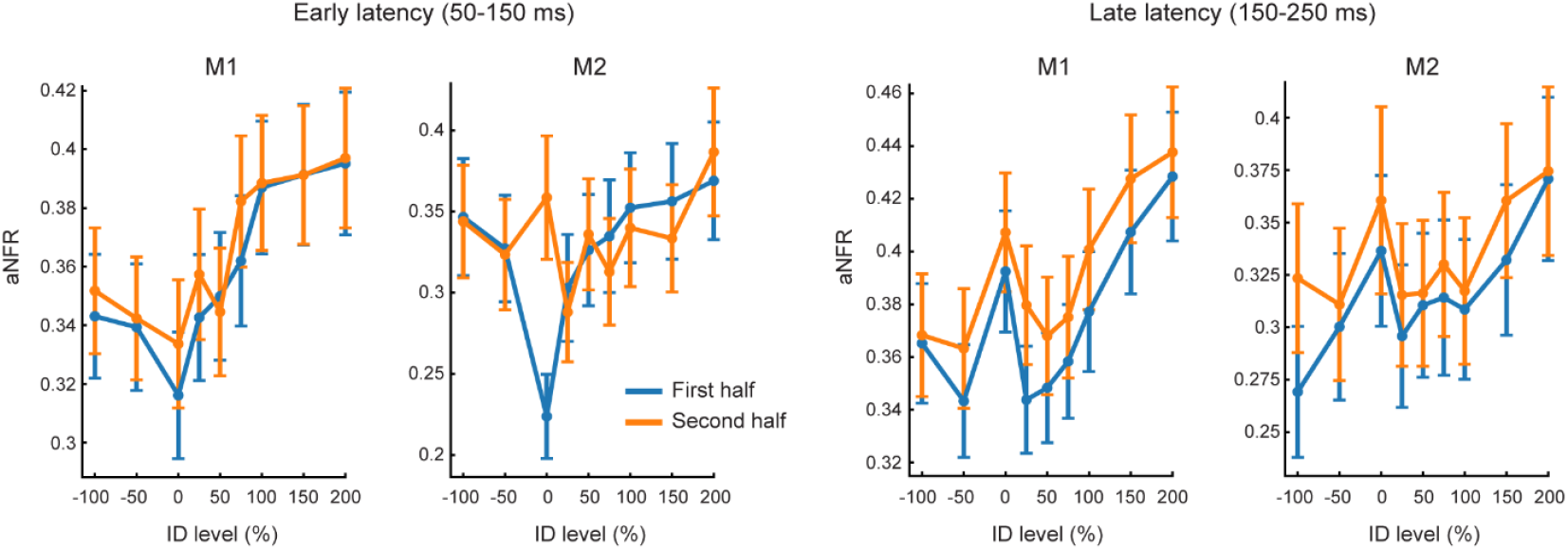
Within session adaptation analysis. Population tuning curves computed during early (50–150 ms; left panels) and late (150–250 ms; right panels) response latencies for both monkeys. Each curve represents the aNFR across neurons for each identity level, separately for the first and second halves of the session.

**Fig. S4.**
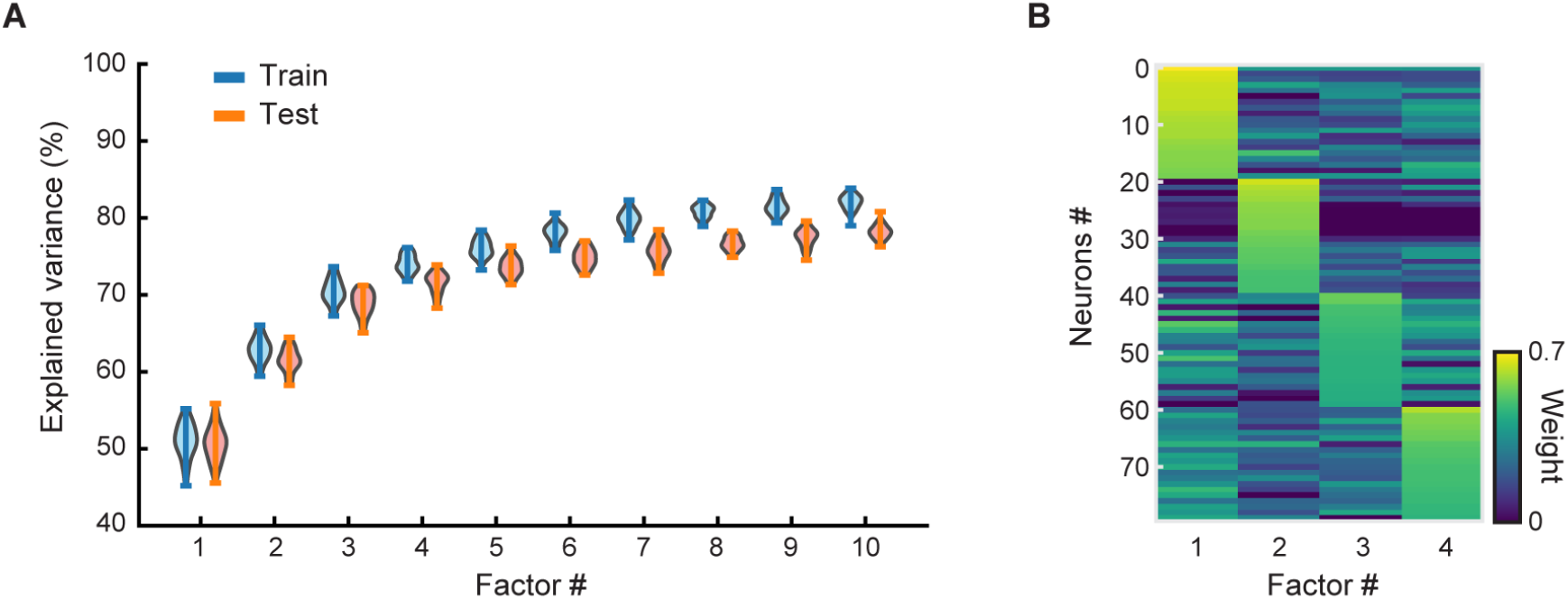
Non-Negative Matrix Factorization performance and neural contributions to factor structure. (**A**) Shuffle-split cross-validation explained variance per number of factors (blue: test set, orange: train set). (**B**) Neuronal weights of the 20 neurons with highest weights on each factor (80 neurons in total).

## Supplementary Audio 1

Stimuli corresponding to the continuum ID1 shown in Fig. S1D.

